# Systematic assessment of sequencing depth requirements for Hi-C-derived metrics

**DOI:** 10.64898/2026.07.28.741297

**Authors:** A. Granovsky, K. Polovnikov

## Abstract

**Background:** Hi-C experiments produce genome-wide chromatin contact maps from which structural features can be quantified across multiple genomic scales, ranging from megabase-scale compartments to kilobase-scale boundaries and chromatin loops. Although high-resolution analyses commonly rely on hundreds of millions of sequenced read pairs, the minimum sequencing depth required for different classes of Hi-C-derived features has not been systematically established. We therefore sought to determine these depth requirements systematically.

**Results:** We performed progressive random subsampling of twelve Hi-C libraries representing multiple cell types and experimental protocols. Loop-density and loop-size inference from the *P(s)* log-derivative was evaluated across all libraries, whereas compartment, insulation, and boundary analyses were performed on a subset of nine libraries with comparable full-depth coverage. Rather than defining sufficient depth as an arbitrary fraction of the full-depth value, we introduced a biologically motivated criterion based on the variability between independent biological replicates. The required sequencing depth for each metric was defined as the point at which subsampling scatter first reached the variability observed between independent biological replicates, representing the accuracy that additional sequencing cannot improve upon. Using this criterion, loop-density estimation reached its biological floor at approximately 10 million read pairs, loop-size estimation at approximately 20 million, insulation scores and boundary detection at approximately 30 million, and compartment eigenvectors at 100-kilobase resolution at approximately 60 million read pairs.

**Conclusions:** Different Hi-C-derived metrics require substantially different sequencing depths to achieve biologically meaningful accuracy. For all metrics, depth-induced variability fell below biological replicate variability well before full sequencing depth was reached. These thresholds provide practical guidance for experimental design and sequencing budget allocation, suggesting that, in many studies, increasing the number of biological replicates is likely to improve reproducibility more effectively than sequencing individual libraries to greater depth.

## Background

Chromosome conformation capture techniques, and Hi-C in particular, have become central tools for studying the three-dimensional organization of the genome (1,2). In a Hi-C experiment, chromatin contacts are converted into a genome-wide matrix in which each entry reports how often two genomic loci are found in spatial proximity. This matrix contains information across a wide range of genomic scales. At megabase scales, it reveals A/B compartments, which reflect the spatial segregation of transcriptionally active and inactive chromatin (1,2). At submegabase scales, it reveals domains and domain boundaries, often quantified through insulation profiles (3). At still finer scales, local enrichments of contacts appear as dots or loops, many of which are associated with cohesin-mediated loop extrusion and convergent CTCF sites (1,4) . Finally, even after averaging over genomic position, the contact probability *P(s)*, where *s* is genomic separation, carries information about the polymer state of chromosomes and about the average properties of cohesin-mediated loops (5,6).

These different features are not extracted from Hi-C maps in the same way. Compartment annotations are usually inferred from eigenvectors of observed/expected contact matrices at relatively coarse resolution. Insulation profiles are local tracks computed from contact density around the diagonal (3) and are therefore more sensitive to the number of contacts in a local genomic window. Individual loops or dots are even more local: they correspond to enriched pixels or small pixel neighbourhoods and are among the most demanding Hi-C features to detect reproducibly. In contrast, the contact probability *P(s)* is a genome-wide average over many pairs of loci at the same genomic separation. Therefore, different Hi-C-derived metrics use different amounts of averaging and should not be expected to have the same dependence on sequencing depth.

Sequencing depth is consequently a central experimental-design parameter. Modern Hi-C libraries are often sequenced to hundreds of millions of read pairs, and the deepest reference maps can contain billions of contacts (1,7,8). Although sequencing throughput has increased and the per-base cost has decreased, sequencing depth remains the main variable cost that scales with the desired number of read pairs. Library preparation introduces a substantial fixed per-sample cost, but deeper sequencing increases the cost of each library linearly. Thus, reducing sequencing depth can substantially lower the sequencing component and make larger designs with more conditions, perturbations or biological replicates more feasible. The practical question is therefore how much sequencing is required for the specific Hi-C-derived metric that a study aims to use.

Several previous studies have addressed related questions, but usually in more restricted settings. One line of work focused on reproducibility of whole contact maps. Methods such as HiCRep, HiC-spector and GenomeDISCO were developed to compare Hi-C maps while accounting for distance dependence, smoothing, spectral properties or diffusion on graphs (9–11). A later systematic comparison showed that reproducibility scores are themselves coverage-dependent and that Hi-C datasets should be compared at matched coverage (12). These studies were essential for quality control and replicate comparison, but their primary object was the reproducibility score, not the depth requirement of individual biophysical features.

A second line of work considered sequencing depth in the context of particular feature callers, especially loop or dot detection. Loop calling is intrinsically local, because a loop is identified as a focal enrichment over its local background. As a result, loop detection is highly sensitive to sequencing depth, resolution, protocol and statistical thresholding. In the systematic comparison of chromosome conformation capture protocols by Akgol Oksuz et al., sequencing depth had a strong effect on the number and strength of detected loops, and formaldehyde-only protocols were more depth-sensitive than protocols using dual FA+DSG cross-linking (7). The same paper further emphasized that compartments can be resolved with tens of millions of valid pairs, whereas individual loop detection may require much deeper sequencing, often hundreds of millions to more than a billion reads depending on the target resolution and protocol (7). Loop-calling tools such as Mustache also used downsampling analyses to test robustness of called loops, but these analyses were necessarily framed around tool-specific loop recovery from very deep reference maps (13). More recently, Parker et al. treated the problem as one of statistical power for differential loop detection and showed that detecting changes in individual loops can require extremely deep maps, with billions of contacts per condition needed for well-powered detection of two-fold loop changes in a large fraction of loops (14).

These studies establish an important point: individual loops or dots are local and demanding Hi-C features. Our study addresses a different question. We do not analyse how individual loop strengths or loop calls degrade upon downsampling. Instead, we focus on genome-wide average metrics of chromosome organization: the mean density and size (genomic length) of random cohesin-mediated loops inferred from the *P(s)* curve, the reproducibility of compartment eigenvectors, the stability of compartment strength, and the stability of genome-wide insulation and boundary tracks. In contrast to the individual loop calls, *P(s)*-derived loop density and loop size are genome-averaged biophysical parameters of loop extrusion, not measurements of a specific CTCF-stabilized loop (a dot) in a contact map. Because they average over many loci, these parameters may remain stable at much lower depth than individual loop calls.

A third limitation of previous depth analyses is the definition of “sufficient” depth. Many studies compare downsampled maps to a full-depth reference and report how a metric changes as a fraction of the full-depth value. This is useful, but it leaves open the question of what fraction is acceptable. Moreover, different metrics also have different natural levels of biological reproducibility, because each Hi-C library samples a finite subpopulation from a heterogeneous ensemble of chromatin conformations rather than a single deterministic genome-folding state. A metric that is nearly identical between independent biological replicates requires very small depth-induced error to be considered reliable. A metric that varies substantially between biological replicates may reach its acceptable accuracy limit at lower sequencing depth, because additional reads cannot remove the biological variability intrinsic to the measurement.

We therefore define a natural threshold for sequencing depth using biological-replicate variability. For each metric, we ask when the scatter introduced by random subsampling becomes comparable to the variability observed between independent biological replicates of the same cell line at full depth. This biological-replicate floor approximates the accuracy limit of the metric under the present experimental conditions: no amount of additional sequencing of one library can eliminate differences that arise between independently prepared cell populations. We also report the corresponding technical-replicate floor, estimated from independent re-sequencing runs of the same library, which represents the tighter reproducibility expected when biological and library-preparation variation are minimized. This framework turns sequencing-depth choice from an arbitrary percentage-of-full-depth rule into a metric-specific comparison against natural biological variability.

Here we apply this framework to twelve publicly available human Hi-C libraries from two studies: eight deeply sequenced libraries from Akgol Oksuz et al., spanning two cell types and two cross-linking protocols, and four in situ Hi-C libraries from Rao et al. (1,7). We subsample raw FASTQ read pairs from full depth down to 200M, 100M, 50M, 25M, 10M and 1M read pairs, and process each subsample through the same pipeline. We then quantify the stability of four classes of metrics: loop density inferred from the short-distance dip of the *P(s)* log-derivative (6), loop size inferred from the loop-size-associated peak (5), compartment eigenvectors, and insulation profiles. The resulting benchmark places these Hi-C-derived metrics on a common scale and provides practical depth recommendations intrinsically tied to biological reproducibility.

## Methods

### Data acquisition and processing

Twelve Hi-C libraries were obtained from SRA: eight from Akgol Oksuz et al. (7) and four from Rao et al. (1). Raw reads were processed with the distiller-nf (15) pipeline using bwa-mem2 (16) alignment with pairtools (17) and cooler (18) to hg38, with mapq30 filtering and multi-resolution .mcool generation from 1 kb to 10 Mb. Contact matrices were balanced (cooler balance) with default mad-max bin filtering. Bins lacking balancing weights (low-coverage or low-mappability bins) carry NaN weights and are excluded from all per-bin correlations. Analyses were restricted to autosomes and chrX.

### Definition of sequencing depth

Throughout, “depth” refers to the number of raw sequenced read pairs – the quantity specified when ordering sequencing – not to mapped, valid, deduplicated, or cis contacts. Subsampling was applied to the raw FASTQ read pairs before alignment, so that each target depth reflects a real sequencing budget; the downstream funnel (mapping rate, valid-pair rate, cis fraction) scales approximately in proportion to raw depth because each subsample is processed independently through the identical pipeline. Metrics are therefore normalised by raw read pairs, and the full single library is the per-library reference.

### Subsampling

Full FASTQ files were subsampled to target depths of 200M, 100M, 50M, 25M, 10M, and 1M read pairs using rasusa software (19). 4–25 independent random subsamples were generated, the count decreasing with depth (25 at 1–50M, to 4 at 200M). Each subsample was independently processed through the full distiller-nf pipeline.

### Loop-density and loop-size inference

Contact probability *P(s)* was computed with cooltools.expected_cis (20) using smoothed (σ = 0.1), genome-aggregated cis expected values on balanced counts. The log-derivative was evaluated numerically from *P(s)*; the short-scale minimum (dip) and the loop-size maximum (peak) were both extracted (Fig. 1): the dip as the min of the slope within a 6–90 kb window, and the peak as the max within a 50–220 kb window. The loop period T (inverse loop density) and effective fragment length *v*_0_*^eff^* were inferred by fitting the dip coordinates (x_min_, y_min_) by the analytical framework of Polovnikov and Starkov (6) yielding loop density = 1/T.

**Figure 1.**
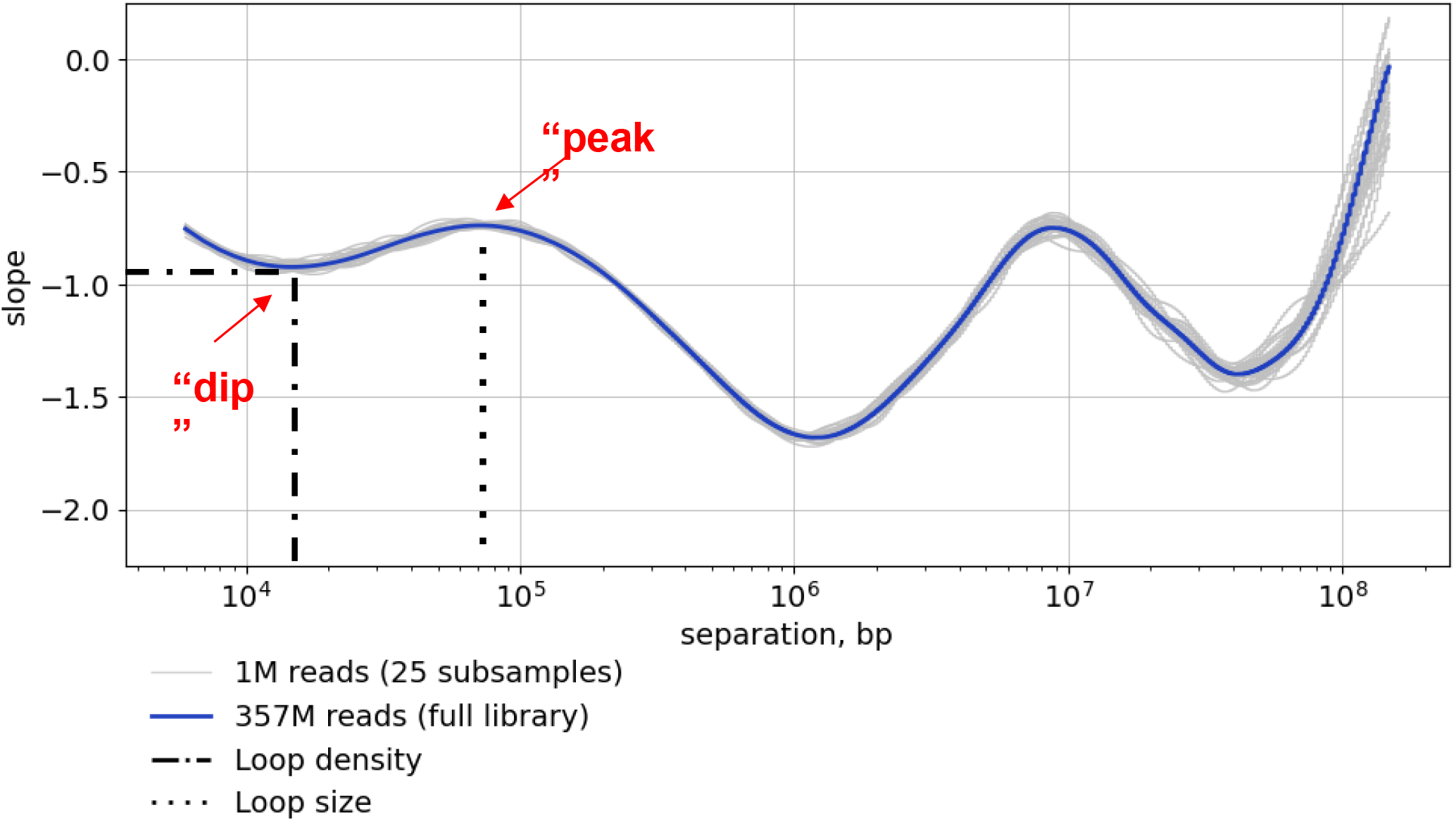
Robustness of *P(s)*-derived loop features to extreme downsampling. Example log-derivative of the contact probability, *d log P*(*s*)/*d log s*, as a function of genomic separation *s* for the hESC-FA-DpnII-R1-T1 Hi-C library. The full-depth library contains 357 million read pairs and is shown in blue; twenty-five of the independent 1M-read subsamples are shown in gray for visual clarity. The short-distance local minimum (“dip”, approximately 15 kb) is used to infer loop density (6), whereas the local maximum (“peak”, approximately 80 kb) is used for the estimate of the loop size (5). Both features remain visually stable after approximately 350-fold downsampling, illustrating the robustness of genome-wide *P(s)*-based metrics.

### Compartment analysis

Compartment eigenvectors were computed at 100 kb resolution using cooltools.eigs_cis (20) with GC-content phasing (hg38 GC track); the first eigenvector E1 was sign-oriented so that GC-rich bins are positive (A compartment). Saddle plots were computed with cooltools.saddle using 38 quantile groups, q_range = [0.025, 0.975]. Compartment strength was defined as (AA + BB) / (AB + BA) using the top/bottom 20% of E1-ranked groups. For each subsample we recorded the Pearson and Spearman correlation of E1 with the full-depth E1 over shared valid bins, the fraction of bins with matching A/B sign, and the scalar compartment strength.

### Insulation and boundary analysis

Insulation scores (3) were computed at 10 kb resolution using cooltools.insulation (20). The 100 kb diamond window is used throughout the main text and Li’s thresholding (21) for boundary calling. Boundary precision, recall, and F1 were computed relative to the full-depth boundary set with ±1,2,3,5 bin tolerance. Boundaries were matched per chromosome by nearest neighbour: a test boundary is a true positive if a full-depth boundary lies within ±х bins, and each reference boundary is counted at most once, so multiple calls falling inside one tolerance window cannot inflate the match count.

### Statistical analysis and threshold estimation

Depth thresholds were defined relative to replicate noise floors. Two replicate levels were used. Biological replicates were independently prepared libraries of the same cell type and protocol, whereas technical replicates were independent sequencing runs of the same library. Variability floors were computed separately for biological and technical replicates using full-depth data. For metrics based on pairwise comparisons between tracks, such as the correlation coefficient and boundary precision, all pairwise comparisons were performed between the replicate libraries belonging to the same dataset, and the resulting values were averaged to obtain a single value for that dataset. Boundary precision was evaluated at each positional tolerance (±1, 2, 3 and 5 bins), and replicate floors were calculated separately for each reported tolerance. For metrics returning a single value per replicate, such as loop size and density, compartment strength, boundary count, and mean boundary strength, the coefficient of variation (CV) of these metrics were computed for each dataset as the standard deviation (SD) divided by the mean across the replicates belonging to that dataset. The per-dataset variability values were then averaged across the datasets. The depth requirement of a metric was defined as the depth at which the subsample variability matches the biological-replicate floor. When single-value metrics are plotted as percentages of their full-depth values, the corresponding floor is shown as a band of 100% ± the averaged CV.

The CV of loop density and loop size was computed within each library. At every depth the mean and the sample SD were taken across the independent subsamples drawn at that depth from one library, and the coefficient of variation was their ratio, *CV = SD / mean*. The value reported against depth is the mean of these per-library CV across the libraries included in the analysis and the shaded band is the SD across the libraries. The full-depth point was set to zero, one full-depth measurement existing per library.

Pearson correlations for the E1 eigenvector and for the insulation track were computed between each subsample of each depth and the full-depth file of the same library. For that the two tracks were inner-joined on bin coordinates (chromosome, start, end), bins with a missing value in either track were discarded, and the coefficient was computed over the remaining bins. Insulation correlations were computed separately for the 50 kb and the 100 kb window. Because multiple independent downsampling replicates were generated at each depth, the correlation was averaged within the library at each depth, then mean and SD across libraries were computed.

Threshold depths were estimated by log-linear interpolation, since sequencing depth was sampled at discrete values and is shown on a logarithmic scale. For each metric, the threshold was defined by the closest pair of consecutive depths whose mean values lay on opposite sides of the biological floor. If *x_k_* and *x_k+1_* denote these depths and y_k_ and y_k+1_ the corresponding metric values, the crossing depth *x* was obtained by interpolation 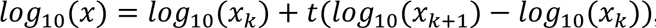, where 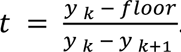. The same interpolation was applied to the upper and lower edges of the SD band around the mean curve and the two depths obtained are reported as the interval within which the crossing lies.

## Results

### Datasets and subsampling strategy

We analysed twelve Hi-C libraries drawn from two studies (Table 1). Eight libraries are from Oksuz et al. (7): four from human embryonic stem cells (H1-hESC) and four from human foreskin fibroblasts (HFFc6), prepared with either a single formaldehyde cross-linker (FA) or the dual DSG+FA cross-linker protocol, with either DpnII alone or a DdeI+DpnII dual digest. The remaining four libraries are from four Rao et al. (1) cell lines (HMEC, IMR90, K562, KBM7), all prepared with *in situ* Hi-C.

**Table 1.** Hi-C libraries used for the sequencing-depth benchmark. The dataset collection comprises twelve publicly available human Hi-C libraries spanning multiple cell lines, chromatin conformation capture protocols, restriction-enzyme strategies, and full-library sequencing depths. All libraries were included in the genome-wide *P(s)*-based analyses of loop-density and loop-size-associated features. Libraries marked with superscript 1 were additionally included in compartment, insulation-score and boundary analyses, for which comparable full-depth coverage was required for cross-library aggregation.

| Dataset | SRA run<br>accessions | Cell line | Protocol | Digestion | Depth,<br>read pairs |
| --- | --- | --- | --- | --- | --- |
| hESC-DSG-DpnII-R1-T1<sup>1</sup> | SRR13601502 | H1-hESC | FA+DSG; DpnII | DpnII | 366M |
| hESC-DSG-DpnII-R2-T1<sup>1</sup> | SRR13601511 | H1-hESC | FA+DSG; DpnII | DpnII | 357M |
| hESC-FA-DpnII-R1-T1<sup>1</sup> | SRR13601520 | H1-hESC | FA; DpnII | DpnII | 357M |
| hESC-FA-DpnII-R1-T2<sup>1</sup> | SRR13601529 | H1-hESC | FA; DpnII | DpnII | 353M |
| HFFc6-DSG-DdeI-DpnII-R1-T1<sup>1</sup> | SRR13601574 | HFFc6 | Hi-C 3.0 | DdeI+DpnII | 361M |
| HFFc6-DSG-DdeI-DpnII-R3-T1<sup>1</sup> | SRR13601583 | HFFc6 | Hi-C 3.0 | DdeI+DpnII | 368M |
| HFFc6-DSG-DpnII-R1-T1<sup>1</sup> | SRR13601592 | HFFc6 | FA+DSG; DpnII | DpnII | 366M |
| HFFc6-DSG-DpnII-R2-T1<sup>1</sup> | SRR13601599 | HFFc6 | FA+DSG; DpnII | DpnII | 352M |
| HMEC | SRR1658680 | HMEC | in situ Hi-C | MboI | 456M |
| IMR90 | SRR1658676 | IMR90 | in situ Hi-C | MboI | 240M |
| K562 | SRR1658694 | K562 | in situ Hi-C | MboI | 591M |
| KBM7<sup>1</sup> | SRR1658708 | KBM7 | in situ Hi-C | MboI | 388M |

The loop-density and loop-size analyses use all twelve libraries, because these metrics are derived from the genome-wide-averaged *P(s)* and their full-depth reference is internal to each library. For the compartment, insulation, and boundaries analyses, only nine libraries were retained: the eight Oksuz et al. libraries plus KBM7, because these metrics are normalised to a per-library full-depth value, and a meaningful cross-library aggregate requires comparable full depths (350-390M). The three excluded Rao cell lines have full-depth read counts of 240M (IMR90), 456M (HMEC), and 591M (K562), all outside the range of the Oksuz libraries.

For each library, the full FASTQ files were subsampled to six target depths: 200M, 100M, 50M, 25M, 10M, and 1M read pairs. At each depth we generated several independent random subsamples to quantify stochastic variability; the number of subsamples was set by the library size, decreasing from 25 at the shallowest targets (1–50M) to as few as 4 at 200 M. The full single library serves as the reference.

### Stability of cohesin-mediated loop size and loop density across sequencing depths

The contact probability *P(s)* is the mean contact frequency between loci separated by genomic distance *s*, averaged over the genome. Across the genomic range typically probed by Hi-C, *P(s)* decays approximately as a power law, reflecting the fractal, crumpled organization of interphase chromatin (2,22,23). At very large *s* close to 10-100Mb, it flattens as loci become confined within chromosome territories (23), whereas at very short *s* about 1-10kb, it is shaped by the finite physical resolution of the Hi-C protocol, including cross-linking efficiency and restriction-fragment size (6). At intermediate scales, chromosome statistics are largely described by a crumpled polymer folded into cohesin-mediated loops by active loop extrusion (5,24,25). Local deviations from scale-free behaviour, reflected as undulations in the log-derivative *d log P*(*s*)/*d log s*, therefore provide a sensitive readout of loop-associated chromatin architecture.

Two such deviations are central to this study: a local minimum (“dip”) at separations about 10–40 kb, and a local maximum (“peak”) at about 100 kb (Fig. 1). The dip arises from the interplay between the loop density and the finite Hi-C capture radius, so its position and depth jointly encode loop density (Fig. S1) (6). The peak, by contrast, reflects the characteristic loop size itself, largely independent of how densely loops are packed (5). Because the two features respond to different structural parameters, tracking them separately allows us to ask whether subsampling degrades the inference of loop density and of loop size at the same rate, or not. The log-derivative of *P(s)* was computed for each subsample and both the short-scale dip and the loop-size peak were extracted (Fig. 1). For each dataset, Tables S1, S2 report the full-depth loop parameters together with the subsample-to-subsample scatter of the loop density as depth decreases.

Before examining the subsampling effects, we first establish two natural noise floors against which subsampling-induced scatter can be compared: the variability between independent biological replicates of the same condition, and the variability between independent technical replicates (Fig. 2A,B; Table S3). Biological replicates are independent Hi-C libraries prepared from separate cell populations of the same line, whereas technical replicates are independent re-sequencing runs of the same library. The former therefore capture both biological and experimental variation, the latter only sequencing noise. Measured at full library depth, the mean coefficient of variation (CV, the ratio of the standard deviation to the mean) across replicates is 13.4% (biological) and 2.5% (technical) for loop density, and 6.5% (biological) and 1.0% (technical) for loop size (Table S3).

**Figure 2.**
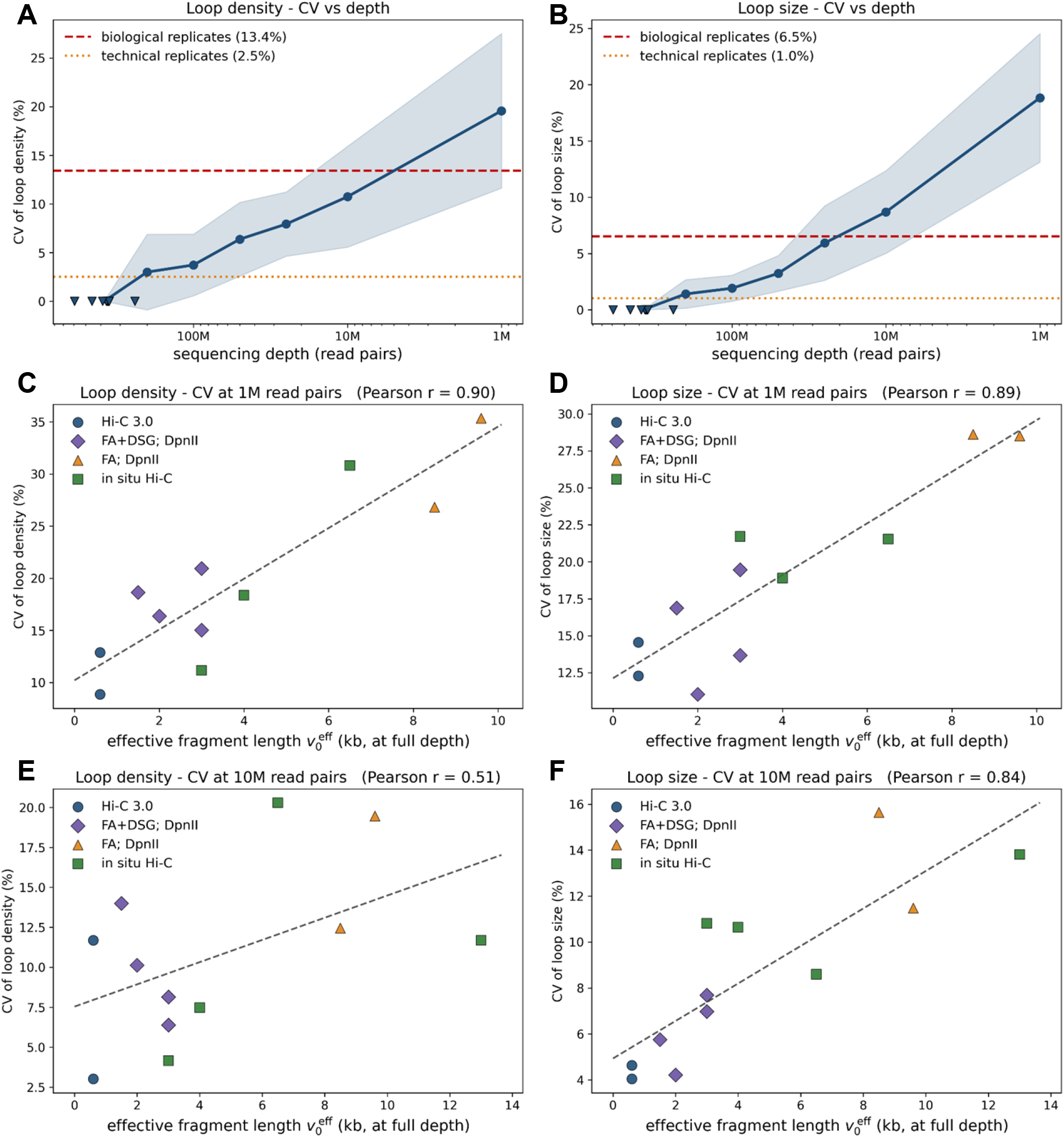
Stability of loop density and loop size under subsampling. Left column: loop density (inferred from the “dip”). Right column: loop size (“peak” position). **(A, B)** Coefficient of variation (CV) of loop density and loop size versus sequencing depth. For each library, CV was computed across independent subsamples drawn at a given depth, the curve shows the mean of these per-library CVs and the shaded band indicates SD across libraries. Dashed lines mark the biological-replicate (13.4% density, 6.5% loop size) and technical-replicate (2.5% density, 1.0% loop size) CV floors, computed as the mean CV across replicate groups at full depth. Inverted triangles along the x-axis mark the original sequencing depth of each library. **(C, D)** The same subsample CV at 1M read pairs, resolved per library, versus the effective fragment length 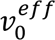 (inferred from the dip at full depth); each point is one library, coloured by protocol, dashed line – least-squares fit (Pearson r = 0.9 for density and 0.89 for loop size). **(E, F)** The same at 10M read pairs (Pearson r = 0.51 and 0.84, respectively). Libraries with smaller 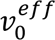 give more stable low-depth estimates.

We find that for all twelve datasets the mean loop density and mean loop size remain close to its full-depth value (± 10%) down to 1-10 million reads (Table S1, S2). Importantly, for both metrics, sequencing depth primarily affects the variation across the subsamples rather than the mean. To place these trends in context, we compare the subsampling-induced scatter to the replicate noise floors established above (Fig. 2A,B). For loop density, the subsampling CV grows from approximately 3% at 200M to approximately 6.4% at 50M and approximately 19.5% at 1M reads. It crosses the technical-replicate floor (2.5%) at 200M reads and reaches the biological-replicate floor (13.4%) at a mean threshold of 5M (range: 1-16M) reads, below which a single subsampled library is noisier than independent biological replicates at the full depth. For loop size, the subsampling CV crosses the biological-replicate floor (6.5%) at mean of 20.6M (range: 6.5-38.4M) reads. The two metrics differ, however, in their relationship to the technical-replicate floor: for density, the subsampling CV falls within the technical replicate floor between 50M and 100M reads, so that subsampling a single deep library reproduces loop density about as tightly as re-sequencing it. For loop size, the subsampling CV never falls to the technical-replicate floor at any depth tested: even at 200M reads the subsample-to-subsample CV (approximately 1.7%) is close to the CV between re-sequencing runs of the same library (1.0%), meaning that re-sequencing reproduces the loop-size estimate more tightly than any level of subsampling can. In practical terms, loop-density estimates from a single library remain within the biological floor down to approximately 5M reads, where the subsampling uncertainty (approximately 0.9 loops/Mb on average across libraries) is comparable to, and does not exceed, the spread between independent biological replicates of the same cell line (0.8–1.6 loops/Mb across the Rao datasets; Table S3). Loop size is somewhat more demanding: its subsampling CV reaches the biological floor at 20M reads, and only at 1M does the scatter (10–25 kb, Table S3) become large relative to the approximately 35 kb spread of loop sizes observed between cell types. Below these depths the subsampling variation exceeds the natural biological variability for both metrics, and a single subsample is a noisier estimate of the underlying parameter than an independent biological replicate sequenced to full depth.

Interestingly, the subsampling uncertainty of both loop-density and loop-size estimates is strongly affected by the experimental protocol, which can be summarized in the effective capture radius at which two loci are registered as a contact in the experiment (6). We express this radius in genomic units as the effective fragment length 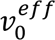, which can be inferred directly from the full-depth dip position (Fig. 2C-F). Across libraries, 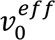 predicts low-depth robustness more strongly than cell type: protocols with tighter capture radius (such as Hi-C 3.0), corresponding to smaller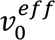, yield more stable estimates under subsampling. Consistently, the HFFc6 DdeI+DpnII libraries, with 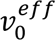, show the smallest scatter at every tested depth, remaining below 0.7 loops/Mb even at 1M read pairs. In contrast, the FA-crosslinked hESC libraries and KBM7 with 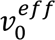, show the largest uncertainty, reaching up to approximately 3.05 loops/Mb at 1M read pairs.

### Compartment eigenvectors and strength

Next we turn to the question of compartment stability with respect to downsampling. A/B compartments were computed at 100 kb resolution across the nine libraries with comparable full depth. The metrics are summarised in Figure 3 (per-depth values in Table S4). We find that the compartment eigenvector is relatively preserved down to approximately 50M read pairs (E1 Pearson r = 93% of full library, Fig. 3A), begins to degrade substantially at 25M (83% correlation), and approaches random expectation at 1M read pairs (correlation near zero, sign agreement about 51%).

**Figure 3.**
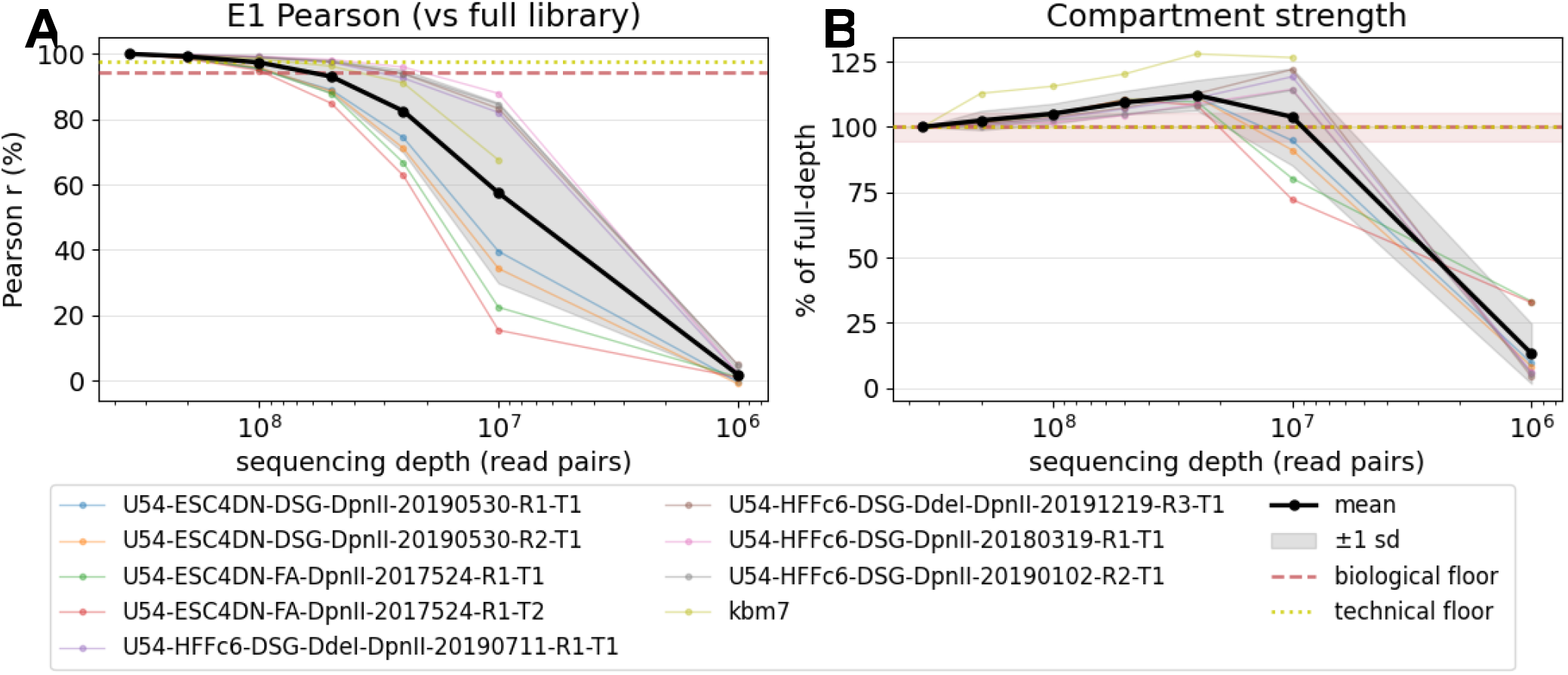
Compartment metrics as a function of sequencing depth (9 libraries). Each panel shows a metric vs subsampling depth. The bold black line is the cross-library mean, the gray band indicates ± 1 s.d. Red and yellow horizontal lines mark the biological- and technical-replicate floors, respectively. (A) E1 Pearson correlation with the full-depth eigenvector. (B) Compartment strength as % of full, showing the non-monotonic inflation at intermediate depths before collapse at 1M.

**Figure 4.**
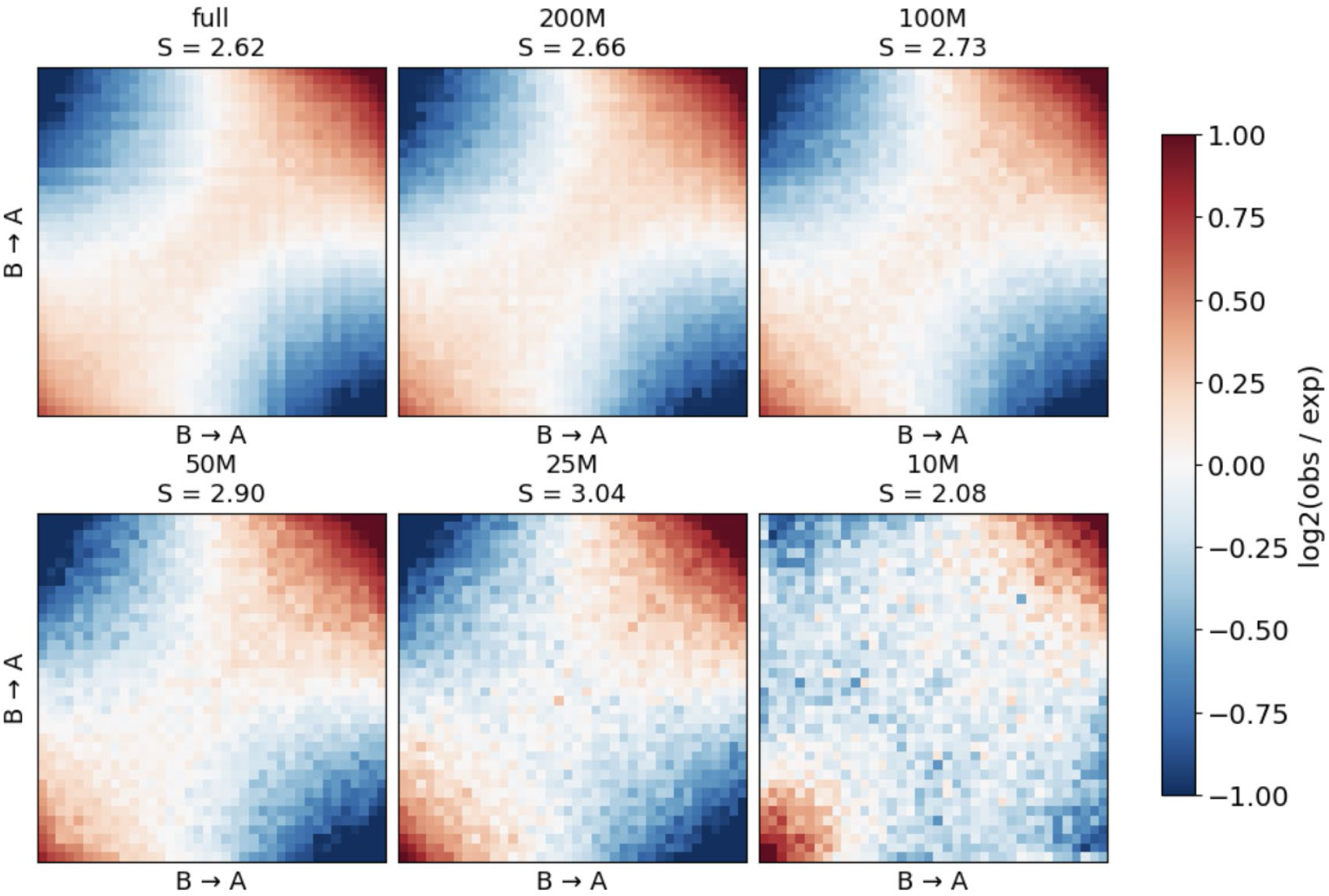
Saddle plots of observed/expected contact enrichment ranked by E1 eigenvector, for one ESC dataset (DSG-DpnII) at five sequencing depths (Full library, 200M, 100M, 50M, 25M, 10M). Red – enriched cis-compartment contacts; blue – depleted trans-compartment contacts.

To place these depth-dependent changes in the context of some natural sample variability, we computed pairwise E1 correlations between independent biological replicates (5-8 replicates from 5 used Rao datasets) and between technical replicates (7–9 re-sequenced runs per library, all from 8 used Oksuz datasets). The mean E1 Pearson r equals 94% between biological replicates and 98% between technical replicates, revealing a significant consistency between the annotations across the replicates. The subsampled E1 Pearson crosses the biological-replicate floor at mean of 61.3M (range: 21.2 - 90.3M) read pairs: below this depth the depth-induced noise exceeds the biological variability (Fig. 3A). The technical-replicate floor is crossed even earlier, at approximately 150M. These estimates reveal a much higher sequencing depth required for the robust compartmental annotation than in the case of loop characteristics analysed above.

Interestingly, compartment strength exhibits an unusual behaviour (Fig. 3B, 4): it is initially slowly rising with downsampling (from 100% to approximately 111% of full at 25M) before dropping to approximately 13% at 1M. This slight initial inflation is a statistical peculiarity of the downsampling procedure that removes numerous weaker, confounding signals such as loops and finer subcompartmental structures, before erasing the stronger E1 compartment signal; this dilution artificially increases the apparent compartment strength as those competing patterns are gradually removed. The biological CV of compartment strength across replicate groups was 6.1% (shown by a red stripe in Fig. 3B), the technical CV was only 0.4% (green stripe).

The aggregate value across the datasets, however, hides an important cell-type dependency: at 10M reads the four hESC libraries decline, down to 84% of full, whereas the five non-hESC libraries (HFFc6 libraries and KBM7) reach 119% of the full (Fig. 3B). All nine nonetheless collapse together at 1M. This is the clearest cell-type-dependent difference we observed; protocol does not produce a comparable separation: the four H1-hESC libraries, including both FA and FA+DSG crosslinking strategies, cluster together in the lower part of the plot, whereas the HFFc6 libraries likewise cluster together above the cell-type-average (Fig. 3B, Table S4).

### Insulation score and boundary calling

To probe the stability of Hi-C domains across sequencing depths, genome-wide insulation scores and domain boundaries were computed at 10 kb resolution and 100kb window. These boundaries were not filtered to any specific size so they demarcated both topologically-associated domains and compartmental/subcompartmental domains. For each subsample, we evaluated Pearson correlation of the insulation track with the full-depth reference, as well as the relative number of called boundaries, relative mean boundary strength, and relative boundary recall (Fig. 5, per-depth values in Table S5).

**Figure 5.**
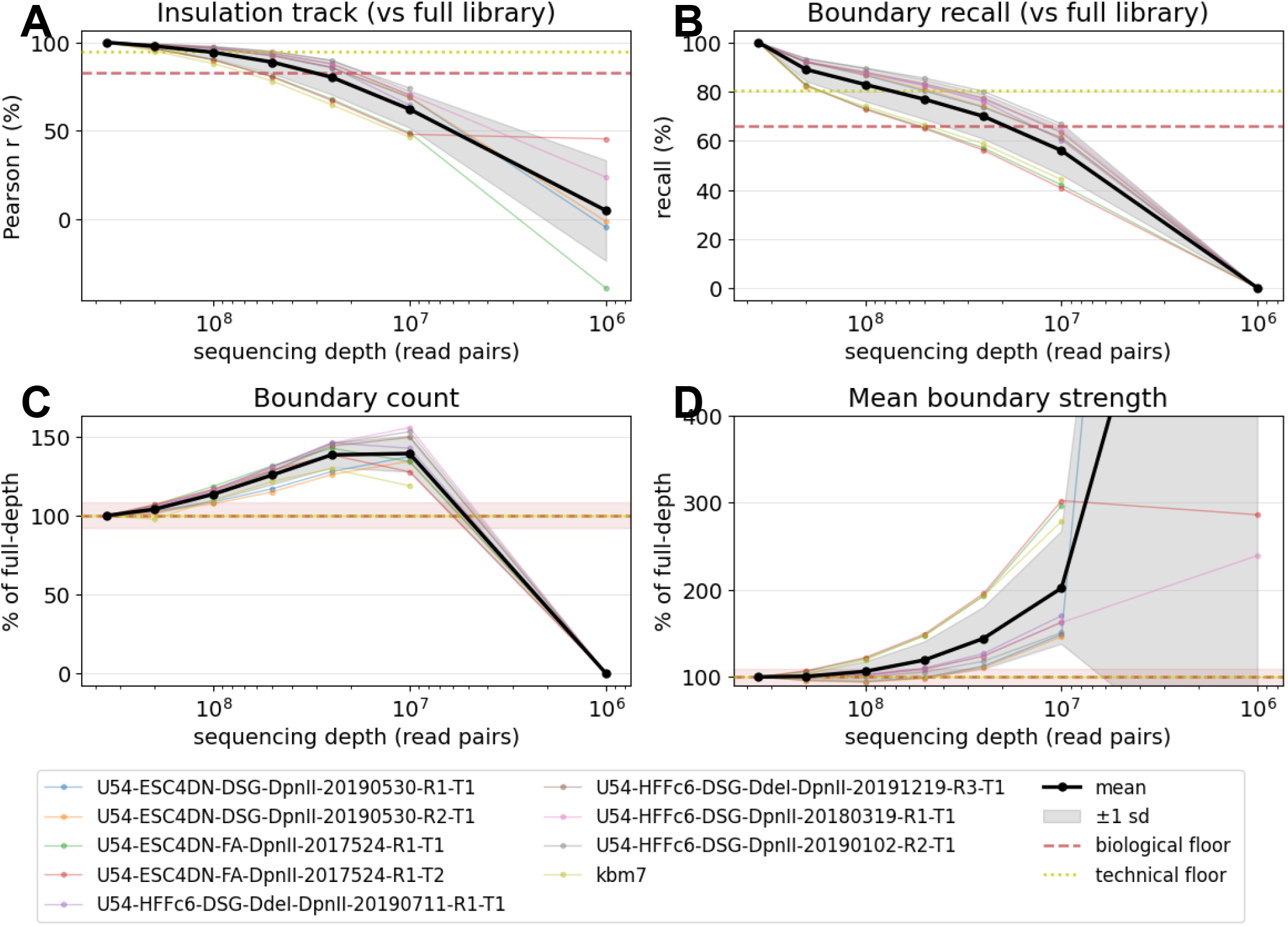
Insulation and boundary metrics as a function of sequencing depth. (n = 9 libraries, 100 kb window, 10 kb resolution). Thin lines: individual libraries; bold: mean ± 1 s.d. Red and yellow lines: biological- and technical-replicate floors. (A) Pearson correlation of the insulation track with the full-depth track. (B) Boundary recall vs the full-depth boundary set (±2-bin tolerance). (C) Number of called boundaries and (D) mean boundary strength as % of full, both showing non-monotonic inflation at intermediate depths.

Similarly to E1 correlation (Fig. 3A), insulation-score correlation degrades substantially already at high sequencing depths: 98% remained at 200M, 89% at 50M, and 62% at 10M. At 1M reads the correlation is virtually zero. In contrast to compartments, the biological-replicate floor for insulation-score correlation is rather low: the mean Pearson r between biological replicates at full depth was only 0.83 (for the technical – 0.97). This points to the intrinsic stochasticity of the domains which are dominated by submegabase topologically-associated domains, while more persistent compartmental boundaries (Fig. 3A) are a minor part of the total. The subsampling curve crosses the low biological floor at mean 30.3M (range 16.3 to 53.0M) reads, which is 2-fold less than 61.3M reads critical depth for the compartments.

Furthermore, the number of called boundaries (Fig. 5C) and mean boundary strength (Fig. 5D) show a non-monotonic inflation pattern similar to compartment strength: boundary count rises to 140% of full at 25M as at intermediate depths noise creates spurious insulation dips that pass the threshold and inflate the count. Mean boundary strength inflates in parallel, rising from approximately 100% of full depth at 200M to 145% at 25M and 200% at 10M (Fig. 5D), before the metric loses meaning at 1M, where no boundaries are called (Fig. 5C, Table S5). The inflation is a direct consequence of the sparser matrix: as coverage falls, the insulation profile becomes noisier and the depth of each local minimum is exaggerated, so the boundaries that still pass the threshold are scored as systematically stronger than in the full-depth reference. The effect is amplified at smaller diamond windows, where fewer pixels enter each insulation value: with a 50 kb window, insulation-track correlation and boundary recall follow trends similar to those observed with the 100 kb window (Fig. S2A,B), whereas mean boundary strength increases substantially, reaching approximately 300% of the full-depth value at 10M reads (Fig. S2D), likely for the same reasons underlying the inflation of compartment strength (Fig. 3B).

At 1M reads the insulation track becomes essentially flat and no boundaries are called. The net effect is captured by boundary recall, which at ±2-bin tolerance drops from 90% at 200M to approximately 56% at 10M (Fig. 5B), with the biological replicate level around 25M. Relaxing the positional tolerance confirms this interpretation: mean recall and both floor estimates improve progressively with wider tolerance (Fig. S3). However, at ±5 bins the tolerance window (50 kb) approaches the size of a small TAD, making the metric less sensitive to biologically meaningful boundary displacements.

### Sequencing cost implications

To translate depth thresholds into real budgets, we note that a library at depth d (in million read pairs) sequenced at 2×150 bp generates approximately 0.3·d Gb of raw sequence (2 × 150 bp × d × 10⁶). Hi-C libraries are compatible with any Illumina paired-end instrument. We price the tiers below assuming a representative $3 per Gb for 2×150 bp reads on the NovaSeq X. (Table 2)(26,27).

**Table 2.** Estimated Illumina sequencing costs per metric tier (2×150 bp paired-end reads). Depth estimates correspond to the biological-replicate floor of each metric, i.e. the depth below which subsampling noise exceeds the variability between independent biological replicates. Sequencing beyond this depth improves precision that is not distinguishable from biological variability. Library preparation is not included.

| Target metric | Reads (M) | Raw Gb | Est. cost |
| --- | --- | --- | --- |
| Loop density | 10 | 3 | >\$9 |
| Loop size | 20 | 6 | >\$18 |
| Insulation | 30 | 9 | >\$27 |
| Compartments | 60 | 18 | >\$54 |
| Standard Hi-C | >350 | >105 | >\$315 |

We recommend at least 10M read pairs for loop-density and 20M loop-size inference from a single library, roughly 0.05x the depth of a full library and therefore about 0.05x its sequencing cost. At these depths, both the density CV and the loop-size CV remain within or close to the biological-replicate floor, so further depth yields diminishing returns relative to the natural variability between independent cell populations.

## Discussion

Since the introduction of Hi-C, sequencing depth has increased by orders of magnitude. The first Hi-C maps, generated from fewer than 10 million informative read pairs, were already sufficient to reveal the large-scale A/B compartmentalization of the genome (2), whereas later high-depth in situ Hi-C maps resolved domains, loops and finer-scale compartmental structure (1). Modern bulk Hi-C libraries are now routinely sequenced to hundreds of millions of read pairs, producing tens to hundreds of gigabases of raw sequence (7,8,28). This increase in depth has transformed Hi-C from a genome-wide assay that primarily revealed broad folding patterns into a quantitative, high-dimensional readout from which chromatin organization can be analysed across multiple genomic scales.

At the same time, the interpretation of Hi-C maps has become increasingly compressed. Many questions about chromosome folding can now be formulated in terms of a limited number of biophysical metrics, such as parameters of random loops (mean loop size and linear density), compartment eigenvectors, compartment strength, insulation profiles and boundary positions. These metrics reflect different scales of genome organization and rely on different amounts of averaging across the contact map. Compartment annotations capture megabase-scale segregation of active and inactive chromatin (1,2). Insulation profiles quantify local separation between neighbouring genomic domains (3). Loop-extrusion models connect local deviations in the contact probability *P(s)* to loop-scale physical parameters (4–6,24,29,30). In this sense, computational and biophysical analysis of Hi-C performs a form of dimensionality reduction: a large contact matrix is projected onto a small number of meaningful structural readouts.

This motivates the central practical question addressed here: how much sequencing depth is required for each Hi-C-derived metric? This question remains relevant despite the continuing increase in sequencing throughput and reduction in per-base cost (27). Hi-C library preparation introduces a substantial fixed per-sample cost, whereas sequencing depth is the main variable cost that linearly scales with the desired number of read pairs. Therefore, reducing depth does not reduce the total experimental cost proportionally, but it can substantially lower the sequencing component and make larger designs with more conditions, perturbations or biological replicates more feasible (Table 2). Importantly, there is no single universal depth requirement. Sequencing-depth requirements depend not only on the genomic scale of the feature, but also on the extent of genomic averaging, the Hi-C protocol and, most importantly, how reproducible the same metric is between independent biological replicates. Our benchmark therefore avoids defining “sufficient depth” as an arbitrary fraction of the full-depth value. Instead, we use an empirical biological floor: the variability observed between independent biological replicates of the same cell line at full depth. This floor approximates the accuracy limit of the metric under the present experimental conditions, because no amount of additional sequencing can eliminate genuine variation between independently prepared cell populations.

On this criterion, the depth hierarchy emerges (Fig. 6). The *P(s)*-derived loop-density dip is the most robust metric, reaching the biological floor at approximately 1-16M read pairs. The loop-size-associated peak reaches its biological floor at approximately 7-38M read pairs. Insulation profiles reach the corresponding floor at roughly 16–53M read pairs. Interestingly, compartment eigenvectors, despite representing large-scale genome organization, are the most demanding under this criterion, crossing their biological-replicate floor at 21-90M read pairs. Thus, the ordering reflects the combination of two effects: how strongly a metric degrades under subsampling, and how reproducible that metric is between independent full-depth biological replicates.

**Figure 6.**
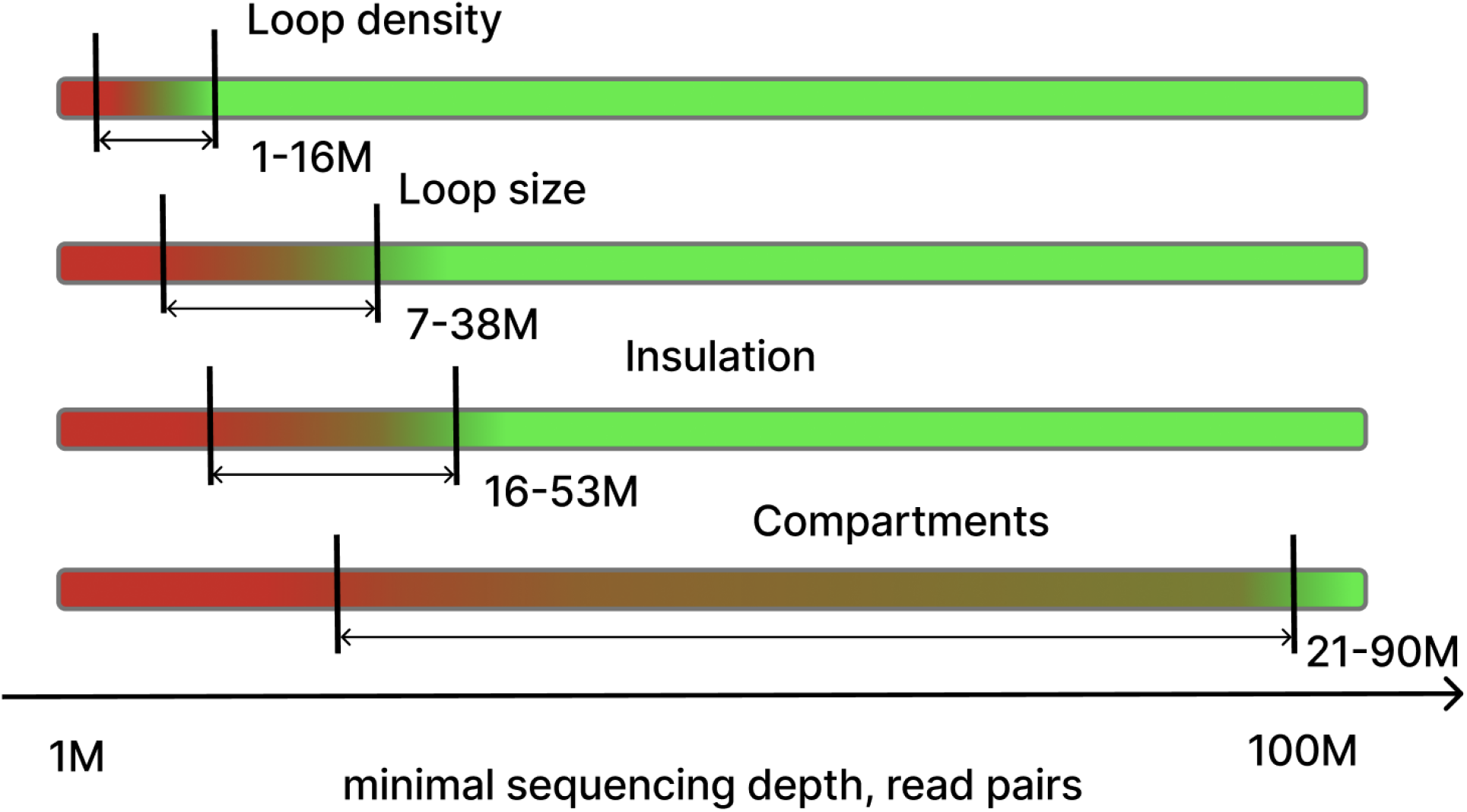
Hierarchy of critical sequencing depths across Hi-C metric families. For loop density and loop size, critical depth was defined as the point at which the mean subsampling CV crossed the corresponding full-depth biological-replicate CV floor. For compartment eigenvectors and insulation profiles, critical depth was defined as the point at which the mean correlation with the full-depth track crossed the corresponding full-depth biological-replicate correlation floor. Central values and ranges were obtained by log-linear interpolation of the mean curve and its ±1 SD envelope across libraries between the two analysed depths bracketing the floor.

A key point is that the mean of many downsampled libraries is not by itself a useful definition of sufficient depth. For the *P(s)*-derived loop metrics, the mean estimate remains close to the full-depth value even after severe downsampling to 1M reads (see Fig. 1, S1; Tables S1, S2). This is expected: averaging over many independent low-depth subsamples effectively reconstructs a deeper experiment. In practice, however, a researcher who deliberately sequences at low depth obtains one random sample of reads, not an ensemble average over many such samples. The relevant question is therefore how far a single low-depth library can deviate from the full-depth estimate. This is why subsample-to-subsample scatter, rather than the mean alone, provides the appropriate measure of reliability.

The loop-density and loop-size stability analysis in Fig. 2 illustrate this distinction. Across all twelve libraries, the mean loop density and mean loop size remain within approximately 10% of their full-depth values down to 1-10M reads. Sequencing depth primarily affects the scatter around these means. Loop density is a global steady-state parameter: in the extrusion picture, it is determined by the total number of chromatin-bound cohesins per genomic length. Its subsampling coefficient of variation increases from approximately 3% at 200M reads to 3.7% at 50M and 19.5% at 1M reads. It crosses the technical re-sequencing floor of 2.5% at 200M, and reaches the biological-replicate floor of 13.4% at approximately 5M reads. Loop size is different: it is an emergent length scale shaped by cohesin processivity together with the distribution and kinetic properties of barriers to loop extrusion (31). Its biological floor is lower, 6.5%, indicating that the inferred loop-size peak is more stable across independent biological replicates than the inferred loop density. This stronger biological reproducibility leaves less room for sequencing-induced scatter, and the biological floor is therefore reached only at approximately 20M reads; the technical floor of 1.0% is not clearly reached at any tested depth. Thus, loop density can be estimated from relatively shallow libraries, whereas loop size requires somewhat deeper sequencing to remain within its narrower natural biological variability.

The robustness of loop-scale inference also depends strongly on experimental protocol. The relevant parameter is the effective fragment length 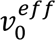, which expresses the Hi-C capture radius in genomic units and can be inferred from the full-depth dip position (6). Libraries with smaller 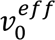 have sharper dip features and tolerate downsampling better (Fig. 2C,E). In our data, the HFFc6 DdeI+DpnII libraries, with 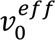 close to 0.6 kb, show the smallest loop-density scatter at every tested depth, remaining below 0.7 loops/Mb even at 1M reads. In contrast, broader-capture libraries, including FA-crosslinked hESC libraries and in situ Hi-C libraries such as KBM7 and IMR90, have larger effective fragment lengths, up to approximately 13 kb, and show substantially greater uncertainty, reaching approximately 3.05 loops/Mb at 1M reads. Similar conclusions were reached in (7), where the number of detected loops in the data was found to be more sensitive to the amount of sequenced reads for protocols with a single cross-linking (FA) than in protocols with double cross-linking (FA+DSG). Protocol choice and sequencing budget are therefore coupled: tighter capture makes low-depth *P(s)*-based inference more stable, whereas broader capture requires deeper sequencing to reach the same precision. Note that in this study the critical read depths are provided as the averages across all available protocols (see Table 1).

The result that compartment eigenvectors require the deepest sequencing under our criterion may appear surprising. A/B compartments were discovered in the first Hi-C study using far fewer reads than modern maps (2). The apparent contradiction comes from the difference between detecting the existence of compartments and quantitatively reproducing a high-resolution compartment track. The broad plaid-like compartment pattern, and even the sign of the E1 eigenvector, can be visibly non-random at low depth. This is sufficient to discover compartmentalization or classify the genome into broad A/B domains. Our benchmark asks a stricter question: at what depth does a 100 kb E1 eigenvector from one subsample reproduce its full-depth counterpart as well as independent full-depth biological replicates reproduce each other? Because the biological-replicate E1 correlation is rather high, approximately r=0.94, depth-induced noise must be very small before it becomes comparable to the biological floor. Thus, compartments can be qualitatively detected at low depth, but stable quantitative compartment annotation requires deeper sequencing.

This comparison also clarifies why insulation and boundary metrics reach their biological floor at lower depth than compartments, despite being more local and more visibly sensitive to coverage. Insulation-track correlation decays substantially under downsampling, falling to approximately 89% at 50M reads and 62% at 10M reads (Fig. 5A). However, the full-depth biological-replicate floor for insulation is much lower than for compartments, approximately r=0.83. In other words, local domain structure is intrinsically less reproducible between independent biological libraries than the compartment eigenvector. Because the biological floor is lower, the subsampling curve reaches it later, at around 30M reads. The minimal useful depth of a metric is therefore not simply the level at which it captures a substantial fraction of the full-depth effect, but the depth at which sequencing noise becomes small relative to the biological variability of that metric.

Scalar summary statistics require special caution. Compartment strength, boundary count and mean boundary strength can appear to improve under downsampling even while the underlying contact map degrades (Fig. 3-5; Fig. S2). These increases should not be interpreted as stronger segregation. Rather, downsampling acts partly as an uncontrolled denoising or coarse-graining procedure: weaker and more local features, including shallow insulation minima, dots, loops and fine subcompartmental patterns, are progressively lost before the strongest large-scale signal disappears. As these competing structures are removed, compartment contrast can appear sharper, while sparse and noisy insulation profiles can generate deeper local minima that pass the boundary-calling threshold. This interpretation is consistent with the broader protocol-dependent trend reported by Akgol Oksuz et al. (7), where protocols that enhance local chromatin features, including loops, tend to show reduced apparent compartment strength.

A simple genome-scale consideration helps interpret why depths near 10–25M read pairs appear as natural lower limits for the most robust metrics. A haploid human genome contains approximately 3 billion base pairs. Twenty million single-end 150 bp reads correspond to about 3 billion sequenced bases, i.e. roughly one haploid genome-equivalent of raw sequence. Since our depths are reported as paired-end read pairs, this corresponds to approximately 10M read pairs for 2×150 bp sequencing. This is not a true coverage threshold for Hi-C contacts: Hi-C reads are not uniformly distributed along the genome, contact maps sample pairs of loci rather than individual bases, and many bins are filtered due to mappability or balancing failures. Nevertheless, it provides useful intuition.

Below roughly 10M read pairs, the experiment contains less than one genome-equivalent of raw sequence, so sparse sampling of genomic regions becomes unavoidable. This raises a related and more fundamental point: the biological-replicate floor may itself reflect finite sampling of the chromatin conformational ensemble. Two independent populations of the same cell line are not identical realizations of one deterministic 3D genome; rather, each Hi-C library averages over a finite number of cells, loci and transient folding states. Similarly to sampling-induced variation during the downsampling procedure, biological replicate variability may therefore be interpreted, at least in part, as sampling of the ensemble of chromosome conformations available to that cell type.

The cost estimates make these exercises practical. With 2×150 bp paired-end sequencing, a library at depth d million read pairs generates approximately 0.3d Gb of raw sequence. At a representative NovaSeq X reagent cost of approximately $3/Gb (26,27), the biological-floor depths correspond to lower-bound sequencing reagent costs of approximately $9 for loop density at 10M read pairs, $18 for loop size at 20M read pairs, $27 for insulation at 30M read pairs and $54 for compartment eigenvectors at 60M read pairs. A standard >350M-read Hi-C library would require more than 105 Gb of raw sequence and more than $315 in sequencing reagents alone. These values exclude library preparation, indexing, quality control, labour and facility overheads, and should therefore be interpreted as lower-bound sequencing costs rather than full experimental costs. Nevertheless, the relative conclusion is robust: several Hi-C-derived metrics reach their biological-floor depth far below the depth of modern full-scale Hi-C libraries, so metric-specific depth selection can substantially reduce the sequencing component of the experiment and make larger replicate-or perturbation-based designs more feasible.

Several limitations bound the scope of these recommendations. First, our benchmark is restricted to bulk Hi-C from human cell lines mapped to the human genome. The numerical thresholds should not be assumed to transfer directly to other genomes, especially genomes with different size, repeat content, mappability, chromosome number or restriction-fragment distribution. Second, as discussed above, the biological floors were estimated from full-depth libraries and represent a lower bound on biological-replicate variability under low sequencing noise; at reduced depth, variability between biological replicates would increase because each replicate would also accumulate independent sampling noise.

Third, random subsampling of deep libraries models the loss of read depth but not the loss of library complexity. Truly shallow experiments may have additional variance from finite unique molecules, PCR duplication, ligation efficiency and other preparation-specific effects that read-level subsampling cannot reproduce. Fourth, the benchmark covers standard bulk Hi-C only; we did not directly test Micro-C (32,33), capture Hi-C (34) single-cell Hi-C (35–37) or other 3C-derived assays. Fifth, our critical-depth thresholds are averaged across protocols, although both our results and previous work (6,7) indicate that cross-linking and restriction strategies affect the effective capture radius and therefore the robustness of *P(s)*-derived metrics to downsampling. Finally, insulation and boundary thresholds depend on caller settings, including bin size, diamond-window size, thresholding method and positional tolerance. For these reasons, the qualitative hierarchy in Fig. 6 is likely more robust than the exact numerical thresholds, which should be interpreted as approximate metric-specific depth ranges rather than universal constants.

## Conclusion

Sequencing-depth requirements for Hi-C are metric-specific and depend on both genomic averaging and biological reproducibility. Using replicate-to-replicate variability at full depth as an empirical biological floor, we find a clear hierarchy: genome-wide *P(s)*-derived loop density is the most robust metric, loop-size inference requires somewhat deeper sequencing, insulation and boundary metrics require intermediate depth, and 100 kb compartment eigenvectors require the deepest sequencing because of their high biological-replicate reproducibility. These results argue against a single universal Hi-C depth standard and provide a practical framework for choosing sequencing depth according to the downstream metric, experimental protocol and intrinsic biological variability.

## List of abbreviations

A/B compartments: Active/Inactive chromatin compartments
P(s): Contact probability as a function of genomic separation *s*
CTCF: CCCTC-binding factor
FA: Formaldehyde
DSG: Disuccinimidyl glutarate
GC: Guanine–cytosine
E1: First eigenvector
CV: Coefficient of variation
TAD: Topologically associated domain
PCR: Polymerase chain reaction

## Declarations

### Ethics approval and consent to participate

Not applicable. This study reanalysed publicly available sequencing datasets and did not generate new human or animal data.

## Consent for publication

Not applicable.

## Availability of data and materials

The public Hi-C datasets analysed in this study are available from the NCBI Sequence Read Archive under the accession identifiers listed in Table 1. The source code and analysis scripts are available at https://zenodo.org/records/21498562 and mirrored at https://github.com/agranovskiii/sequencing-depth-requirements. Additional processed summary tables used to generate the figures are included in the Supplementary Information.

## Authors’ contributions

A.G. performed data processing, subsampling analyses and figure generation. K.P. designed the study, supervised the analysis and contributed to interpretation. A.G. and K.P. wrote and revised the manuscript. Both authors read and approved the final manuscript.

## Acknowledgements

Not applicable

## Supplementary

**Figure S1.**
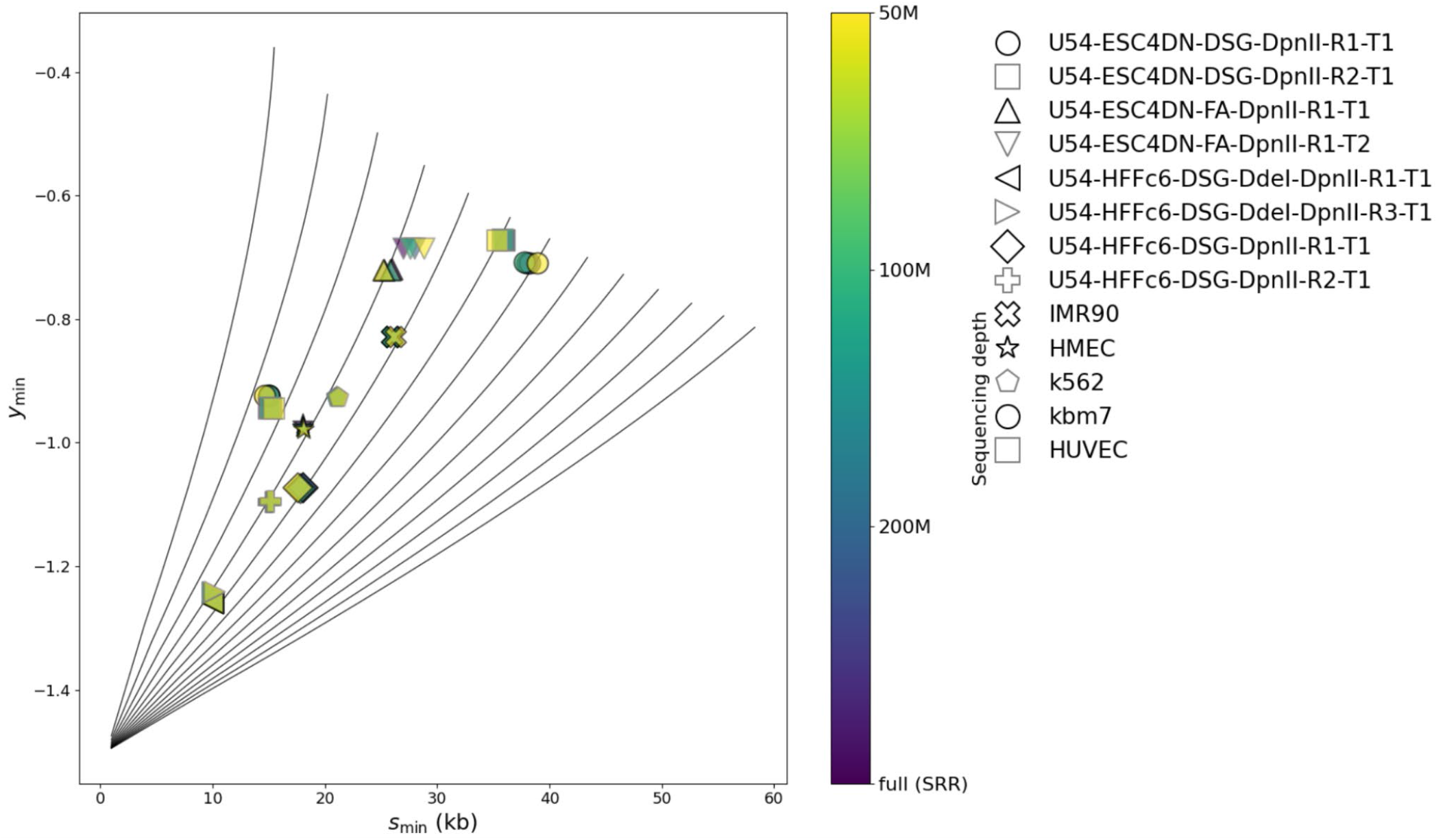
Fountain diagram of fitted *P(s)* derivative dip coordinates. Gray curves: analytically predicted positions for the inverse cohesin density T = 60–300 kb (step 10 kb). Filled markers: experimentally fitted dip positions extracted from *P(s)* derivative. Colour encodes sequencing depth; marker shape identifies the dataset.

**Figure S2.**
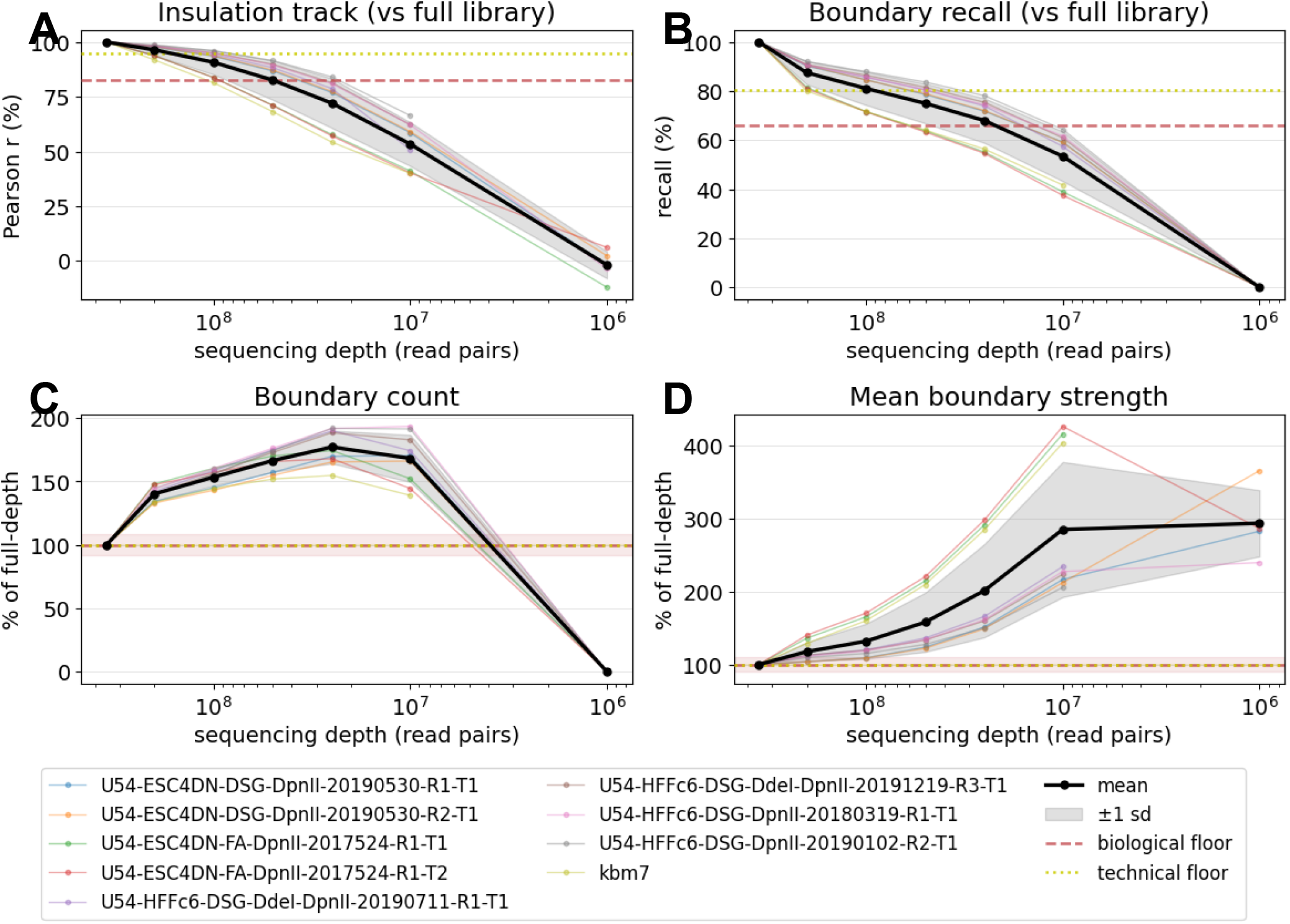
Insulation metrics as a function of sequencing depth. (n = 9 libraries, 50 kb window, 10 kb resolution). Thin lines: individual libraries; bold: mean ± 1 s.d. Red and yellow lines: biological- and technical-replicate floors. (A) Pearson correlation of the insulation track with the full-depth track. (B) Boundary recall vs the full-depth boundary set (±2-bin tolerance). (C) Number of called boundaries and (D) mean boundary strength as % of full, both showing non-monotonic inflation at intermediate depths.

**Figure S3.**
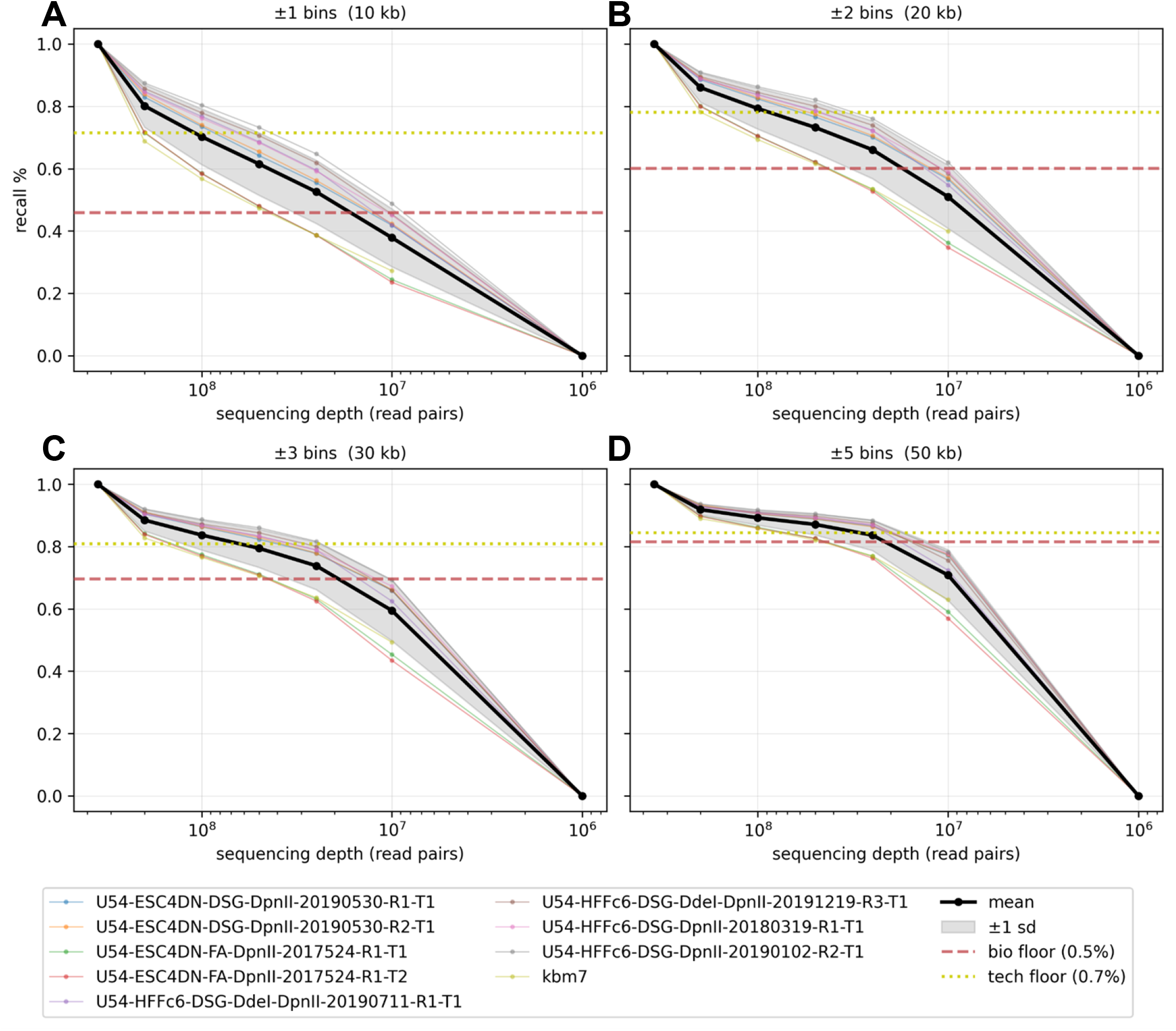
Boundary recall across tolerance levels. Each panel shows boundary recall (fraction of full-depth boundaries recovered) as a function of sequencing depth for a given positional tolerance: (A) ±1 bin (10 kb), (B) ±2 bins (20 kb), (C) ±3 bins (30 kb), (D) ±5 bins (50 kb). Thin lines represent individual libraries; the bold black line is the cross-library mean ± 1 s.d. Dashed red and dotted yellow lines mark the biological and technical floors, respectively. Increasing the tolerance progressively raises recall at every depth.

**Table S1.** Loop density at full depth and at each subsampling depth (mean ± s.d. across independent subsamples, loops/Mb). 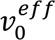 and loop period T shown at full depth. SD values of 0.00 indicate that the standard deviation was non-zero but smaller than the displayed precision after rounding.

| Dataset | Density<br>full<br>(l/Mb) | $v_0^{eff}$<br>(kb) | T<br>(kb) | 200M | 100M | 50M | 10M | 1M |
| --- | --- | --- | --- | --- | --- | --- | --- | --- |
| HMEC | 9.09 | 3 | 110 | $8.71 \pm 0.44$ | $8.64 \pm 0.41$ | $8.33 \pm 0.00$ | $8.56 \pm 0.36$ | $8.52 \pm 0.95$ |
| IMR90 | 7.14 | 6.5 | 140 | $7.14 \pm 0.00$ | $7.36 \pm 0.30$ | $7.25 \pm 0.69$ | $7.36 \pm 1.49$ | $7.32 \pm 2.26$ |
| hESC-DSG-DpnII-R1-T1 | 11.11 | 3 | 90 | $11.11 \pm 0.00$ | $11.11 \pm 0.00$ | $11.67 \pm 0.72$ | $11.47 \pm 0.73$ | $11.41 \pm 2.39$ |
| hESC-DSG-DpnII-R2-T1 | 10.00 | 3 | 100 | $10.00 \pm 0.00$ | $10.00 \pm 0.00$ | $10.22 \pm 0.46$ | $10.16 \pm 0.82$ | $10.49 \pm 1.56$ |
| hESC-FA-DpnII-R1-T1 | 8.33 | 8.5 | 120 | $8.36 \pm 0.57$ | $8.08 \pm 0.35$ | $8.52 \pm 0.70$ | $8.77 \pm 1.09$ | $9.02 \pm 2.42$ |
| hESC-FA-DpnII-R1-T2 | 8.33 | 9.6 | 120 | $7.69 \pm 0.00$ | $7.97 \pm 0.63$ | $7.69 \pm 0.73$ | $8.28 \pm 1.61$ | $8.61 \pm 3.05$ |
| HFFc6-DSG-Ddel-DpnII-R1-T1 | 7.69 | 0.6 | 130 | $7.69 \pm 0.00$ | $7.69 \pm 0.00$ | $7.69 \pm 0.00$ | $7.79 \pm 0.23$ | $7.82 \pm 0.69$ |
| HFFc6-DSG-Ddel-DpnII-R3-T1 | 9.09 | 0.6 | 110 | $9.09 \pm 0.00$ | $9.09 \pm 0.00$ | $8.70 \pm 0.82$ | $8.31 \pm 0.97$ | $7.86 \pm 1.01$ |
| HFFc6-DSG-DpnII-R1-T1 | 7.69 | 2 | 130 | $7.18 \pm 0.59$ | $7.49 \pm 0.46$ | $7.55 \pm 0.51$ | $7.54 \pm 0.76$ | $6.78 \pm 1.11$ |
| HFFc6-DSG-DpnII-R2-T1 | 9.09 | 1.5 | 110 | $8.60 \pm 0.97$ | $8.70 \pm 0.87$ | $8.12 \pm 1.08$ | $8.50 \pm 1.19$ | $8.86 \pm 1.65$ |
| K562 | 8.33 | 4 | 120 | $8.33 \pm 0.00$ | $8.08 \pm 0.35$ | $8.14 \pm 0.31$ | $7.95 \pm 0.59$ | $8.20 \pm 1.47$ |
| KBM7 | 5.56 | 13 | 180 | $5.56 \pm 0.25$ | $5.69 \pm 0.18$ | $5.54 \pm 0.30$ | $5.59 \pm 0.65$ | $6.12 \pm 1.14$ |

**Table S2.** Loop size (peak position, kb) at full depth and at 200M, 100M, 50M, 10M and 1M reads (mean ± s.d. across independent subsamples). Peak slope varies negligibly with depth (s.d. ≤ 0.02 at all tested depths).

| Sample | Loop size<br>full (kb) | Loop size<br>200M | Loop size<br>100M | Loop size<br>50M | Loop size<br>25M | Loop size<br>10M | Loop size<br>1M |
| --- | --- | --- | --- | --- | --- | --- | --- |
| hESC-DSG-DpnII-R1-T1 | 71 | $70.5 \pm 0.6$ | $70.4 \pm 1.5$ | $71.4 \pm 1.8$ | $71.1 \pm 2.7$ | $71.2 \pm 5.0$ | $74.6 \pm 10.2$ |
| hESC-DSG-DpnII-R2-T1 | 75 | $74.0 \pm 0.0$ | $74.2 \pm 0.4$ | $73.8 \pm 1.9$ | $74.2 \pm 3.4$ | $75.0 \pm 5.7$ | $71.9 \pm 13.9$ |
| hESC-FA-DpnII-R1-T1 | 74 | $73.8 \pm 1.3$ | $75.2 \pm 1.5$ | $74.3 \pm 4.4$ | $73.8 \pm 7.6$ | $73.9 \pm 11.6$ | $78.2 \pm 22.4$ |
| hESC-FA-DpnII-R1-T2 | 82 | $82.0 \pm 1.4$ | $80.8 \pm 3.8$ | $81.7 \pm 4.1$ | $82.3 \pm 9.7$ | $84.0 \pm 9.6$ | $86.0 \pm 24.5$ |
| HFFc6-DdeI-DpnII-R1-T1 | 104 | 103.5 ± 0.6 | 103.4 ± 1.1 | 105.0 ± 1.4 | 104.5 ± 2.9 | 104.5 ± 4.2 | 112.6 ± 16.4 |
| HFFc6-DdeI-DpnII-R3-T1 | 102 | 101.2 ± 1.3 | 101.8 ± 0.8 | 101.9 ± 1.2 | 101.2 ± 3.0 | 101.2 ± 4.7 | 100.1 ± 12.3 |
| HFFc6-DSG-DpnII-R1-T1 | 106 | 104.8 ± 1.0 | 105.0 ± 1.6 | 105.1 ± 2.4 | 106.2 ± 2.4 | 105.4 ± 4.4 | 102.9 ± 11.4 |
| HFFc6-DSG-DpnII-R2-T1 | 100 | 99.5 ± 1.0 | 99.0 ± 1.0 | 100.0 ± 2.2 | 99.9 ± 3.8 | 99.7 ± 5.7 | 102.2 ± 17.3 |
| HMEC | 100 | 99.8 ± 1.3 | 98.4 ± 3.6 | 97.9 ± 3.7 | 98.7 ± 7.7 | 101.1 ± 10.9 | 101.8 ± 22.1 |
| IMR90 | 91 | 89.8 ± 0.5 | 90.0 ± 1.9 | 91.2 ± 2.9 | 90.2 ± 4.1 | 88.0 ± 7.6 | 91.7 ± 19.8 |
| K562 | 89 | 88.8 ± 1.7 | 89.4 ± 1.5 | 89.7 ± 2.4 | 88.9 ± 4.7 | 88.7 ± 9.4 | 92.7 ± 17.6 |
| KBM7 | 100 | 99.2 ± 5.2 | 99.0 ± 1.6 | 99.1 ± 6.1 | 102.0 ± 11.5 | 99.4 ± 13.7 | 101.2 ± 16.3 |

**Table S3.** Coefficient of variation (%) of loop density, loop size, and peak slope within biological and technical replicates. Mean biological CV: density 13.4%, peak size 6.5%, slope 4.7%; mean technical CV: density 2.5%, peak size 1.0%, slope 0.09%.

| Group / sample | Replicate type | n | Density CV<br>(%) | Peak size CV<br>(%) | Peak slope CV<br>(%) |
| --- | --- | --- | --- | --- | --- |
| ESC4DN-DSG-DpnII | biological | 2 | 8.71 | 2.38 | 0.86 |
| HFFc6-DSG-DdeI-DpnII | biological | 2 | 8.57 | 1.1 | 2.25 |
| HFFc6-DSG-DpnII | biological | 2 | 12.13 | 3.4 | 4.9 |
| HMEC | biological | 6 | 11.41 | 2.98 | 2.87 |
| IMR90 | biological | 8 | 22.99 | 21.4 | 9.12 |
| K562 | biological | 6 | 18.12 | 6.37 | 10.02 |
| KBM7 | biological | 5 | 12.05 | 7.97 | 3.1 |
| U54-ESC4DN-DSG-DpnII-R1-T1 | technical | 9 | 6 | 1.38 | 0.12 |

| Group / sample | Replicate type | n | Density CV (%) | Peak size CV (%) | Peak slope CV (%) |
| --- | --- | --- | --- | --- | --- |
| U54-ESC4DN-DSG-DpnII-R2-T1 | technical | 9 | 0 | 0.94 | 0.13 |
| U54-HFFc6-DSG-DdeI-DpnII-R1-T1 | technical | 9 | 0 | 0.58 | 0.08 |
| U54-HFFc6-DSG-DdeI-DpnII-R3-T1 | technical | 9 | 0 | 0.98 | 0.1 |
| U54-HFFc6-DSG-DpnII-R1-T1 | technical | 7 | 9.17 | 0.85 | 0.05 |
| U54-HFFc6-DSG-DpnII-R2-T1 | technical | 8 | 0 | 1.42 | 0.08 |

**Table S4.**
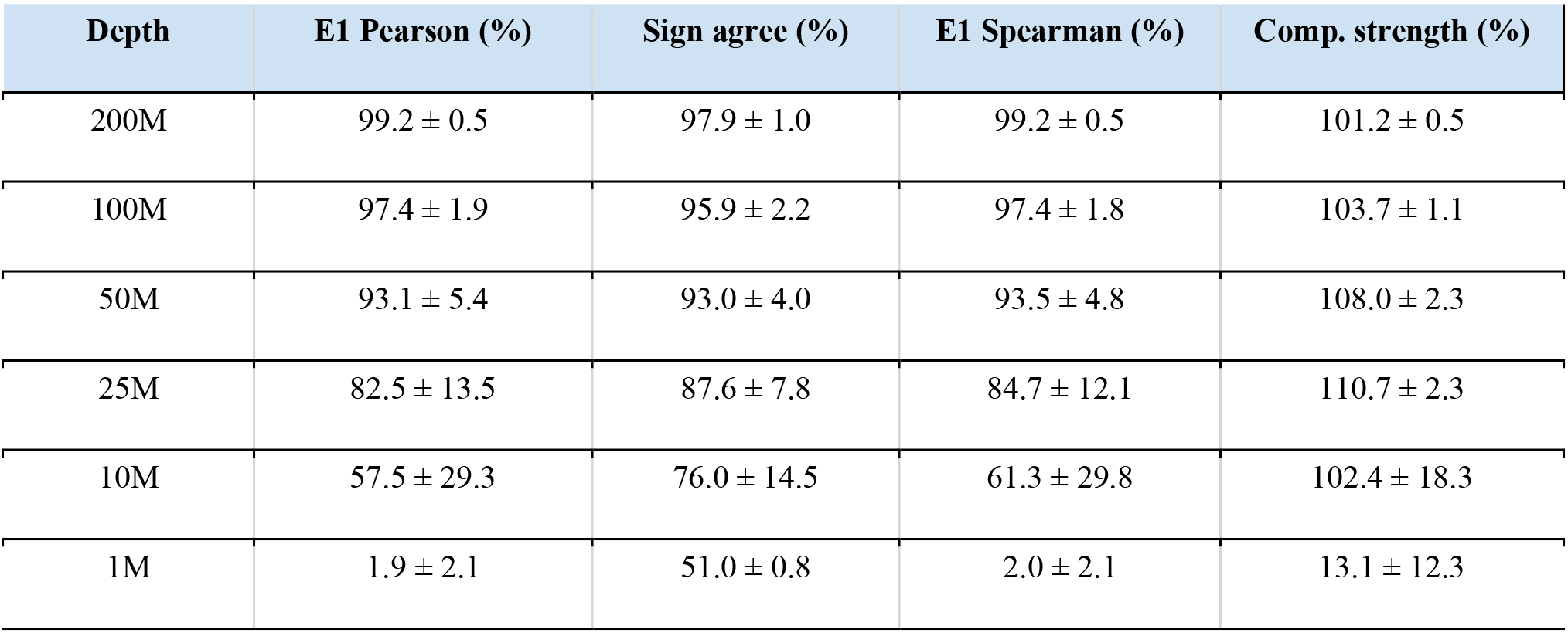
Compartment metrics (% of full-depth value, mean ± s.d. across 9 datasets). E1 eigenvector computed at 100 kb.

**Table S5.** Insulation metrics (% of full, mean ± s.d., 9 datasets; 100 kb window, 10 kb resolution, ±2bin (20kb) tolerance).

| Depth | Ins. Pearson (%) | Boundary F1 (%) | Boundary prec. (%) | Boundary recall (%) | Boundary count (%) | Boundary strength (%) |
| --- | --- | --- | --- | --- | --- | --- |
| 200M | 97.9 ± 1.5 | 86.6 ± 5.8 | 84.4 ± 6.6 | 89.0 ± 5.0 | 105.6 ± 3.1 | 101.5 ± 5.0 |
| 100M | 94.4 ± 3.8 | 77.1 ± 8.1 | 72.1 ± 8.7 | 82.9 ± 7.2 | 115.4 ± 5.6 | 107.2 ± 12.7 |
| 50M | 88.8 ± 6.9 | 67.6 ± 8.6 | 60.5 ± 8.8 | 76.9 ± 8.5 | 127.8 ± 7.6 | 120.4 ± 23.5 |
| 25M | 80.3 ± 10.4 | 58.2 ± 8.4 | 50.0 ± 7.7 | 69.9 ± 9.7 | 140.7 ± 8.4 | 145.5 ± 39.8 |
| 10M | 62.1 ± 11.2 | 46.3 ± 7.5 | 39.6 ± 5.8 | 56.0 ± 10.5 | 141.3 ± 9.6 | 204.0 ± 71.2 |
| 1M | 4.7 ± 31.9 | 0.0 ± 0.0 | 0.0 ± 0.0 | 0.0 ± 0.0 | 0.0 ± 0.0 | 1001.7 ± 1280.3 |

